# The potent broadly neutralizing antibody VIR-3434 controls Hepatitis B and D Virus infection and reduces HBsAg in humanized mice

**DOI:** 10.1101/2022.09.09.507326

**Authors:** Florian A. Lempp, Tassilo Volz, Elisabetta Cameroni, Fabio Benigni, Jiayi Zhou, Laura E. Rosen, Julia Noack, Fabrizia Zatta, Hannah Kaiser, Siro Bianchi, Gloria Lombardo, Stefano Jaconi, Hasan Imam, Leah B. Soriaga, Nadia Passini, David M. Belnap, Andreas Schulze, Marc Lütgehetmann, Amalio Telenti, Andrea L. Cathcart, Gyorgy Snell, Lisa A. Purcell, Christy M. Hebner, Stephan Urban, Maura Dandri, Davide Corti, Michael A. Schmid

## Abstract

**Background & Aims:** Chronic hepatitis B is a major global public health problem, and coinfection with hepatitis delta virus (HDV) worsens disease outcome. Here, we describe a hepatitis B virus (HBV) surface antigen (HBsAg)-targeting monoclonal antibody (mAb) with the potential to promote functional cure of chronic hepatitis B and D to address this unmet medical need.

**Methods:** HBsAg-specific mAbs were isolated from memory B cells of HBV vaccinated individuals. In vitro neutralization was determined against HBV and HDV enveloped with HBsAg representing eight HBV genotypes. Human liver-chimeric mice were treated twice weekly with a candidate mAb starting three weeks post HBV inoculation (spreading phase) or during stable HBV or HBV/HDV coinfection (chronic phase).

**Results:** From a panel of human anti-HBs mAbs, VIR-3434 was selected and engineered for pre-clinical development. VIR-3434 targets a putative conserved, conformational epitope within the antigenic loop of HBsAg and neutralized HBV and HDV infection with >12,000-fold higher potency than Hepatitis B Immunoglobulins in vitro. Neutralization was pan-genotypic against strains representative of HBV genotypes A-H. In the spreading phase of HBV infection in human liver-chimeric mice, a parental mAb of VIR-3434 (HBC34) prevented HBV dissemination and intrahepatic HBV RNA and cccDNA increase. In the chronic phase of HBV infection or co-infection with HDV, HBC34 treatment decreased circulating HBsAg by >1 log and HDV RNA by >2 logs.

**Conclusions:** This in vitro and in vivo characterization identified the potent anti-HBs mAb VIR-3434, which reduces circulating HBsAg and HBV/HDV viremia in human liver-chimeric mice. VIR-3434 is currently in clinical development for treatment of patients with chronic hepatitis B or D.

**Lay summary:** Chronic infection with hepatitis B virus places approximately 290 million individuals worldwide at risk for severe liver disease and cancer. Currently available treatments result in low rates of functional cure or require lifelong therapy that does not eliminate the risk of liver disease. We isolated and characterized a potent, human antibody that neutralizes hepatitis B and D viruses and reduces infection in a mouse model. This antibody could provide a new treatment for patients with chronic hepatitis B and D.

**Highlights:**

- Identification of a human mAb VIR-3434 that potently neutralizes HBV and HDV
- VIR-3434 targets a conserved, conformational epitope of the HBsAg antigenic loop
- VIR-3434 treatment blocks intrahepatic HBV spread in human liver-chimeric mice
- VIR-3434 treatment reduces circulating HBsAg and HDV RNA in co-infected mice
- Data have enabled clinical development of VIR-3434 against chronic hepatitis B/D

## Introduction

Chronic infection with HBV is one of the leading causes of liver cirrhosis and hepatocellular carcinoma. Despite a widely available and efficacious vaccine against HBV, approximately 290 million people are currently living with chronic Hepatitis B (CHB) resulting in an estimated 820,000 deaths per year [1].

Approved treatments for CHB include nucleos(t)ide reverse transcriptase inhibitors (NRTIs) and pegylated-interferon alpha (PEG-IFNα); however, neither treatment results in high rates of HBsAg loss. Therapy with NRTIs leads to elimination of HBV DNA in circulation but has limited effect on HBsAg levels. Long-term NRTI therapy reduces but does not eliminate the risk of hepatocellular carcinoma and requires lifelong treatment for most patients. Treatment with PEG-IFNα can induce HBsAg loss, but only in a small fraction of patients (<10%) and the treatment is generally associated with increased flu-like side effects related to the immunomodulatory effects of the treatment [2]. Additionally, Hepatitis B Immunoglobulins (HBIG), which are polyclonal human anti-HBs antibodies, purified from the serum of vaccinees, have long been used for preventative indications, specifically perinatal mother-to-child HBV transmission and of re-infection after liver transplantation [3]. These limited treatment options underscore the need for novel therapies against CHB that are finite and well tolerated. Effective treatments aim at inducing HBV functional cure, which is defined as sustained suppression of HBV DNA and loss of HBsAg off treatment.

HBV is a small, enveloped virus from the *Hepadnaviridae* family, with a relaxed-circular DNA genome packaged into a capsid formed by dimers of the viral core protein. The capsid is surrounded by a lipid bilayer, into which the three HBV surface antigen proteins (HBsAg) – large (L-HBsAg), middle (M-HBsAg) and small (S-HBsAg) – are embedded. All three surface proteins share the same C-terminal S-HBsAg domain [4]. HBsAg has multiple functions in the viral replication cycle, including facilitating hepatocyte binding and entry by reversible attachment of HBsAg to heparan sulfate proteoglycans (HSPGs) on the surface of hepatocytes [5], followed by non-reversible, high-affinity interaction of the viral preS1 domain of the L-HBsAg with the cellular receptor sodium taurocholate co-transporting polypeptide (NTCP) [6]. HBsAg is also involved in the assembly and secretion of progeny virions and, most importantly, can be secreted in absence of capsid and the HBV genome, forming spherical or filamentous subviral particles (SVPs) that exceed the number of circulating virions by 3-6 orders of magnitude [4]. Such a high level of circulating HBsAg is thought to represent a viral immunotolerance strategy that can exhaust adaptive immune responses [7]. To circumvent immune exhaustion, removing the tolerogenic HBsAg from circulation could help restore T- and B-cell responses and lead to control of HBV infection [8-10]. An ideal therapeutic strategy may combine immune-stimulatory mechanisms with virus neutralization.

HBsAg can be hijacked by HDV, a satellite virus of HBV from the *Kolmioviridae* family with a circular, negative-sense single-strand RNA genome. HDV uses HBsAg for its own envelopment and dissemination and, consequently, the two viruses share the same entry pathway into hepatocytes via NTCP. Therefore, HDV infection occurs only as co- or superinfection with HBV and is associated with the most severe form of viral hepatitis [11, 12]. Here, we characterized a potent and broadly neutralizing mAb that neutralized HBV and HDV infections in vitro, efficiently blocked intrahepatic viral spread and reduced HBsAg as well as HBV and HDV viremia in human liver-chimeric mice.

## Materials and Methods

### Isolation of anti-HBs monoclonal antibodies (mAbs) from human memory B cells

The use of blood cells from healthy human HBV vaccinated donors was approved by the ethical committee “*Comitato Etico Canton Ticino*” (Switzerland). All participants gave written informed consent. Individuals were selected based on high anti-HBs serum antibody titer, as tested by a standard diagnostic method. Total IgG^+^ memory B cells were immortalized from peripheral blood mononuclear cells via a previously described method [13]. After 2 weeks of culture, the B cell supernatants were analyzed for their capacity to bind to HBsAg from three serotypes (adw, adr, ayw) by ELISA. To isolate human mAbs, mRNA from B cells of wells with HBsAg binding was reverse transcribed via RT-PCR, cloned, and produced recombinantly as IgG1 via transient transfection into mammalian cell lines. MAbs were affinity-purified using HiTrap Protein A columns and sterilized via passage through 0.22 µm filters.

**For further details regarding the materials and methods used, please refer to supplementary information and CTAT table**

## Results

### Isolation and engineering of a potent anti-HBs human mAb from a vaccinated donor

To identify broadly neutralizing antibodies against HBV, we isolated IgG^+^ memory B cells from peripheral blood mononuclear cells of HBV vaccinated donors and immortalized them using Epstein-Barr Virus and CpG [13]. The supernatants from B cell cultures were analyzed for binding to HBsAg by ELISA. Selected mAbs were produced as recombinant IgG1 and neutralized HBV more potently than previously characterized mAbs against preS1 (Ma18/7 [14]), against HBsAg (17.1.41, 19.79.5 [15]) or polyclonal HBV immune globulin (HBIG) (**Fig. 1A**). To determine binding behavior, we performed a binding competition assay by ELISA. The original B-cell derived mAb HBC34 showed excellent neutralization capacity and did not compete with the other potently neutralizing mAbs, indicating that it recognizes a distinct functional epitope on HBsAg (**Fig. S1A**). Engineering of the HBC34 Fab light chain variable region for improved developability and introducing two sets of mutations in the Fc generated the preclinical lead VIR-3434 (alias HBC34-dev-LS-GAALIE). The LS Fc mutation (M428L N434S) was introduced to extend in vivo half-life via increased binding to neonatal FcR (FcRn) [16-18] and the GAALIE/XX2 Fc (G236A A330L I332E) to augment effector functions [19-21]. Importantly, these mutations in the Fab and Fc of VIR-3434 did not alter HBsAg binding affinity compared to the parental HBC34 antibody (**Fig. 1B**).

**FIGURE 1.**
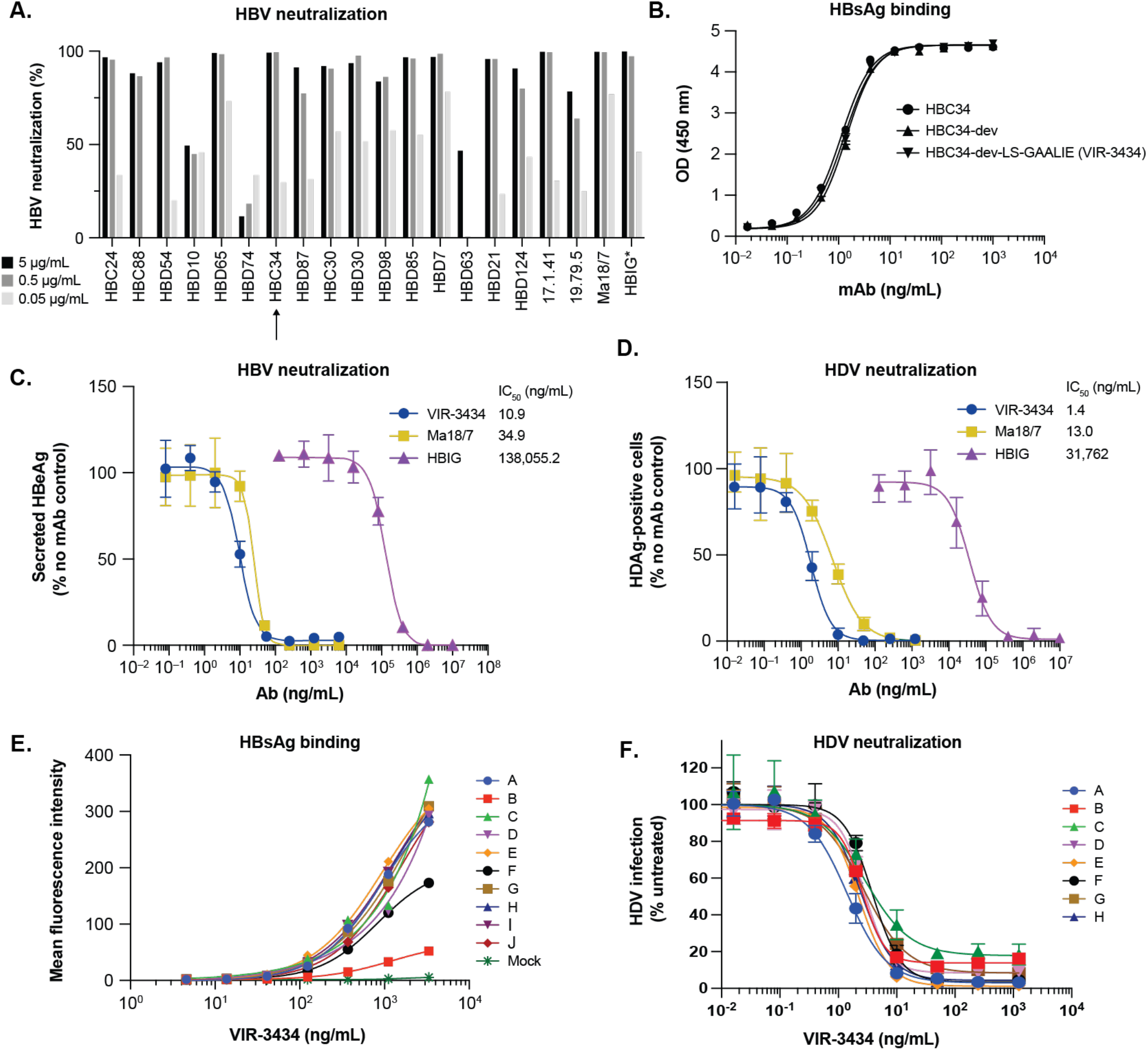
Pan-genotypic neutralization of HBV and HDV infection. (A) HBV neutralization of a panel of human mAbs using HepaRG cells. *Hepatitis B Immune Globulin (HBIG) was tested at 1000x higher concentrations 5,000, 500, and 50 µg/ml. (B) ELISA binding of HBC34 development variants to HBsAg serotype adw. (C) Secreted HBeAg as marker of infection of primary human hepatocytes (PHH) with HBV (genotype D) in the presence of VIR-3434, Ma18/7 mAb or HBIG. (D) Immunostaining for HDAg of Huh7-NTCP cells infected with HDV (genotype 1, enveloped with HBsAg of HBV genotype A) in the presence of respective antibodies. (E) Flow cytometry binding of VIR-3434 to Expi293 cells transfected with HBsAg of genotypes A-J. (F) VIR-3434 neutralizing HDV enveloped with HBsAg of 8 different HBV genotypes on Huh7-NTCP cells.

### VIR-3434 neutralizes HBV and HDV infection with pan-genotypic activity in vitro

To assess the *in vitro* neutralizing activity of VIR-3434 against HBV, primary human hepatocytes (PHH) were infected with HBV in the presence of VIR-3434, Ma18/7 or HBIG (**Figs. 1C** and **S1B**). All three antibodies led to a concentration-dependent reduction in HBeAg (**Fig. 1C**) and HBsAg (**Fig. S1B**) secretion, indicating the inhibition of HBV infection and replication. VIR-3434 had a >12,000x lower IC_50_ value based on HBeAg secretion compared to HBIG (10.9 *vs*. 138,000 ng/ml) and showed an increased potency (>3x) compared to Ma18/7 (34.9 ng/ml), a mAb that binds to the preS1 region of the L-HBsAg (**Fig. 1C**) [14]. In addition, VIR-3434 efficiently neutralized HDV (genotype 1, enveloped with HBsAg of HBV genotype A) in Huh7-NTCP cells with an IC_50_ value of 1.4 ng/ml, while Ma18/7 and HBIG neutralized with IC_50_ values of 13.0 ng/ml (>9x higher) and 31,800 ng/ml (>22,000x higher), respectively (**Fig. 1D**). We next investigated the binding of VIR-3434 to HBsAg from all ten HBV genotypes (A-J) in transiently transfected Expi293 cells. VIR-3434 demonstrated pan-genotypic binding activity, with only genotypes B and F demonstrating lower binding levels (**Fig. 1E**). In this qualitative assay, this may be due to the lower expression levels of HBsAg that was detected with these genotypes. To evaluate the breadth of neutralization of VIR-3434, we generated HDV enveloped with HBsAg from eight different HBV genotypes (A-H). VIR-3434 potently neutralized virus harboring all tested HBsAg genotypes with similar potency (IC50 range 1.4 – 4.2 ng/mL), including genotype B and F (**Fig. 1F**). Taken together, these data show that VIR-3434 potently and broadly neutralizes HBV and HDV *in vitro* with pan-genotypic activity.

### VIR-3434 targets a conserved epitope on the antigenic loop (AGL) of HBsAg

To characterize whether VIR-3434 recognizes a linear or conformational epitope, we analyzed the binding of VIR-3434 to HBsAg preparations produced in yeast or in a human hepatoma cell line (PLC/PRF/5) by Western Blot. VIR-3434 recognized HBsAg only under non-reducing conditions, indicating that VIR-3434 binds to a conformational epitope (**Fig. 2A**). To further map the epitope, we utilized the Chemical Linkage of Peptides onto Scaffolds (CLIPS) technology (Pepscan), which identifies conformational and discontinuous epitopes by constraining antigen-derived peptides to adopt looped structures that mimic the structure of the peptide in context of the full protein [22]. Motif 1 (_114_STTSTGPCRTC_124_) in S-HBsAg was required for VIR-3434 binding (**Fig. 2B**), while a second motif 2 (_145_GNCTCIPIPSSWAF_158_) stabilized binding (**Fig. 2B**). To evaluate conservation of the epitope in circulating HBV strains, 28,331 HBV sequences from the HBV database (HBVdb [23]) were aligned to a reference sequence (GenBank ACU26993.1) using pairwise alignments. Remarkably, >99% conservation was observed for 7 out of 11 amino acids in motif 1 and 12 out of 14 amino acids in motif 2, with high conservation and/or conservative substitutions for the remaining positions (**Fig. 2C, S2A** and **S2B**). Of note, 19 known natural HBV variants or vaccine escape mutants carry mutations within the epitope. VIR-3434 demonstrated binding activity to all but one of the evaluated HBsAg variants by flow cytometry (**Fig. 2D**). However, binding activity to T123N or the vaccine escape variant G145R was decreased, and binding to the T123N/C124R double-mutation was completely abrogated (**Fig. 2D**) [24, 25]. We next generated HDV enveloped with HBsAg harboring the T123N, T123N/C124R or G145R mutations to assess the ability of VIR-3434 to neutralize these mutants. Despite decreased binding to T123N in flow cytometry (**Fig. 2D**), VIR-3434 efficiently neutralized this mutant (**Fig. 2E**). In contrast, the T123N/C124R double mutant was not neutralized at the antibody concentrations tested but also demonstrated a 6-fold reduced viral titer during production of the virus stock compared to WT or the other HBsAg mutants (**Fig. 2F**). The removal of the cysteine at position 124 may destabilize the antigenic loop and possibly result in reduced viral fitness, as observed previously [26]. Additionally, VIR-3434 demonstrated decreased neutralization of the G145R variant. Among the 28,331 sequences available in the HBVdb, the C124R and G145R mutations were only detected at low frequencies of 0.03% and 0.77%, respectively. Overall, VIR-3434 binds to a highly conserved conformational epitope in the HBsAg antigenic loop, resulting in broad pan-genotypic neutralizing activity.

**FIGURE 2.**
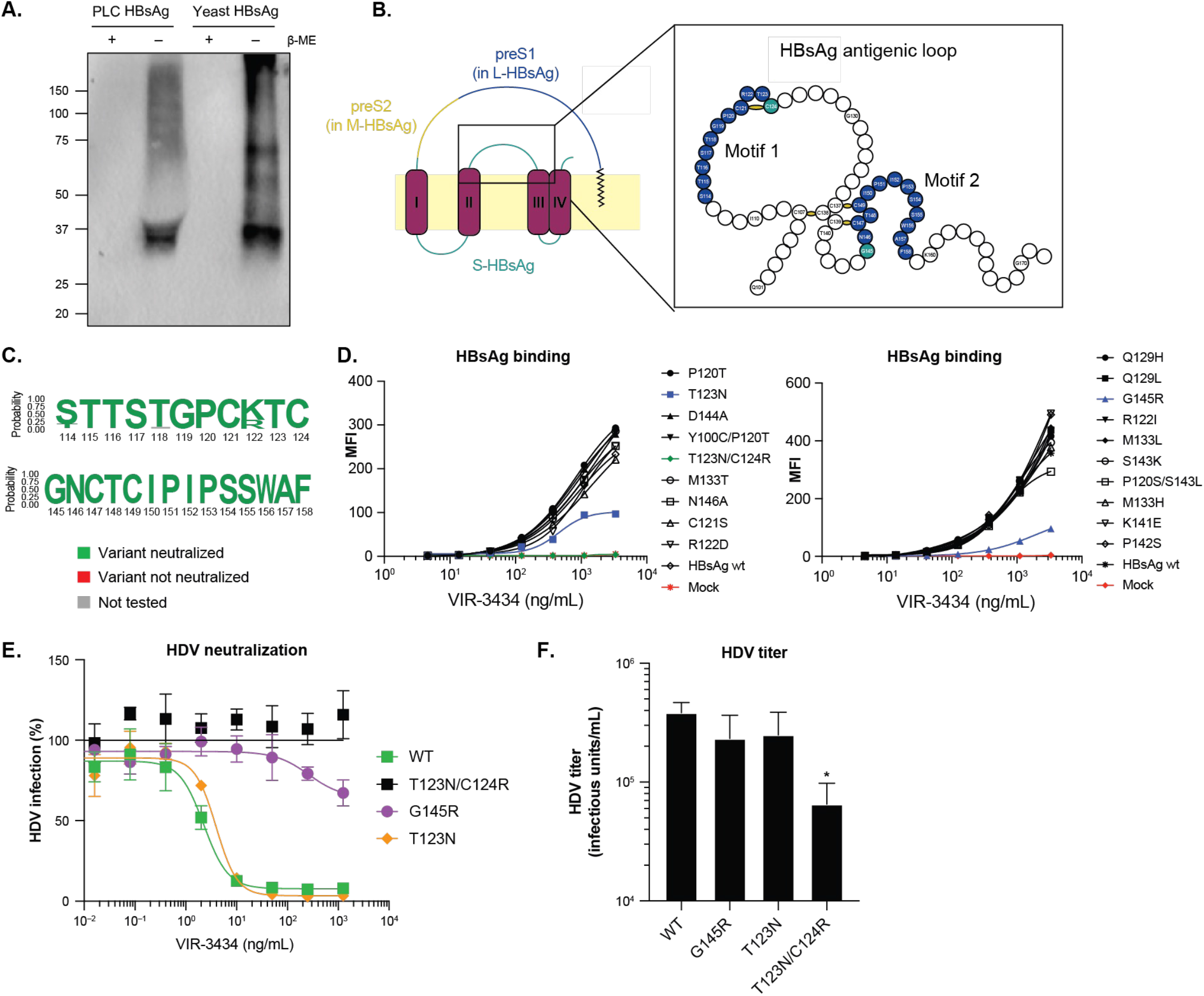
VIR-3434 binds a conserved epitope of the HBsAg antigenic loop. (A) Western Blot with HBsAg separated under non-reducing or reducing conditions and probed with HBC34 for detection. (B) Epitope mapping using a library of linear and looped peptides (CLIPS technology). The VIR-3434 epitope consists of two motives (blue) in the antigenic loop of HBsAg. (C) Epitope conservation plots based on HBV sequence data from HBVdb. Residues of virus variants colored based on VIR-3434 neutralization (green) or loss of neutralization (red); grey variants were not tested. Only frequencies of >2% were visualized. (D) Binding via flow cytometry of VIR-3434 to HBsAg variants transiently expressed in Expi293 cells. (E) Neutralization of HDV enveloped with the respective HBsAg variants. (F) Infectious titer by titration of HDV enveloped with wildtype HBsAg (genotype D) or one of three virus variants on HuH7-NTCP cells. Shown geometric mean ± SD of two independent experiments; statistically significant differences relative to WT by one-way ANOVA (p-value* ≤ 0.05).

### Potential mechanism of VIR-3434-mediated HBV neutralization

A hallmark of CHB is the persistence of high HBsAg levels in the blood. The high excess of HBsAg subviral particles (SVPs) may act as a decoy, competing with antibodies binding to infectious virions. We thus assessed the *in vitro* neutralization capacity of VIR-3434 in the presence of exogenous HBsAg SVPs derived from yeast or PLC cells. Western Blot confirmed that the yeast-derived HBsAg contained only S-HBsAg, whereas PLC cell supernatant contained a mixture of S-, M-& L-HBsAg (**Fig. S3**). Both forms of exogenous HBsAg competed with VIR-3434 neutralization, resulting in a shift (up to 25-fold) of the neutralization IC50 (**Fig. 3A**). At concentrations exceeding 1 µg/ml, competition could be overcome to achieve complete neutralization, even in the presence of 2,000 IU/ml of HBsAg.

**FIGURE 3.**
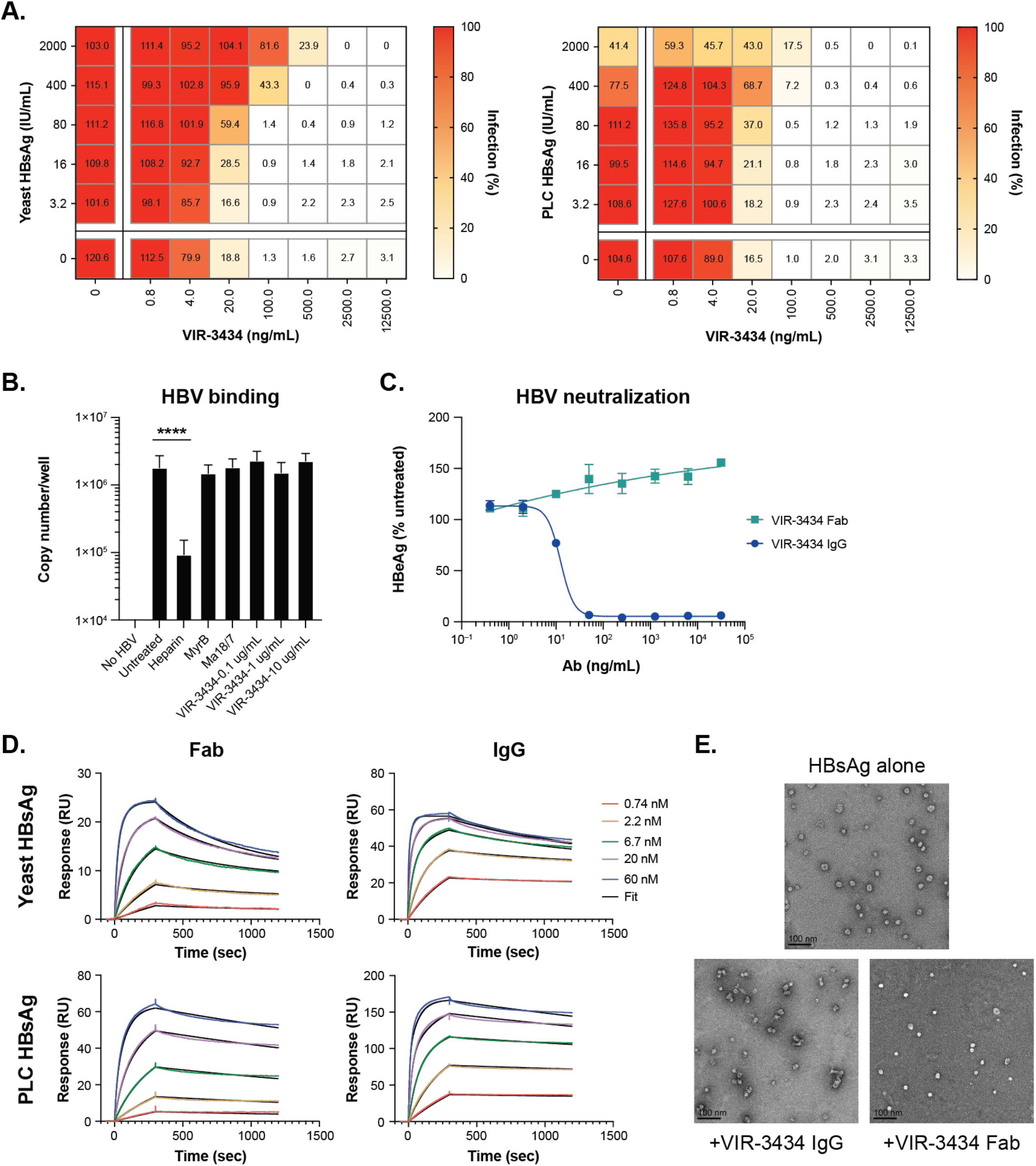
VIR-3434 IgG neutralizes infection and aggregates HBsAg in immune complexes. (A) HBV virus stock (50 IU HBsAg/mL) premixed with indicated concentrations of HBsAg was incubated with VIR-3434 and added to primary human hepatocytes for infection. (B) Binding of HBV to HepG2-NTCP cells in the presence of inhibitors quantified by qPCR. Shown is the mean ± SD of nine data points from a single experiment. Statistical differences were analyzed by one-way ANOVA. p-value **** p ≤ 0.0001. (C) HBV neutralization using VIR-3434 IgG or Fab on primary human hepatocytes. (D) SPR analysis of VIR-3434 IgG or Fab binding to immobilized HBsAg. (E) Yeast-derived HBsAg (1,500 IU/mL) was pre-incubated with 5 µg/mL VIR-3434 IgG or the equivalent molar amount of isolated VIR-3434 Fab fragments and imaged by negative-stain electron microscopy.

Next, we set out to investigate the mechanism of action by which VIR-3434 inhibits HBV infection. The attachment of HBV to hepatocytes is mediated by i) low-affinity binding of HBsAg to HSPGs––cell-surface glycoproteins modified by glycosaminoglycans (GAG) interacting with the preS1 domain and the antigenic loop of HBsAg––and ii) by high-affinity binding of preS1 to the entry receptor NTCP [5]. We exposed HepG2-NTCP cells to HBV in the presence or absence of VIR-3434 and quantified cell-bound virus by quantitative PCR. Neither VIR-3434, the entry inhibitor Myrcludex B [27], nor the preS1-targeting mAb Ma18/7 were able to block binding of HBV to NTCP-expressing cells. Only heparin, a soluble, sulfated GAG could efficiently block attachment of HBV to host cells (**Fig. 3B**), as previously reported [5]. These data indicate that VIR-3434 neutralizes HBV infection not via blocking GAG-mediated attachment but at a later step of infection.

We next compared the neutralization capacity of the full VIR-3434 IgG molecule with that of its Fab fragment. While VIR-3434 IgG potently neutralized HBV, VIR-3434 Fab did not (**Fig. 3C**), suggesting that high-avidity bivalent binding of VIR-3434 IgG to virions may be required to block viral entry. However, surface plasmon resonance (SPR) experiments showed that VIR-3434 IgG and Fab have comparable or only modestly different apparent affinities to immobilized HBsAg (K_D,app_ <1 nM for IgG *vs*. <1-8 nM for Fab) (**Fig. 3D, Suppl. Table S1**). Therefore, an alternate explanation is that neutralization by VIR-3434 requires the cross-linking of virions to form antibody-antigen immune complexes, which is mediated by full-length IgG but not the Fab fragment. To test this hypothesis, we analyzed the morphology of yeast-derived HBsAg in complex with VIR-3434 by negative stain electron microscopy. The untreated HBsAg control as well as HBsAg incubated with the VIR-3434 Fab contained symmetrical, monodispersed, spherical SVPs of ∼20 nm diameter (**Fig. 3E**). In contrast, pre-incubation of HBsAg with VIR-3434 IgG induced cross-linking of SVPs (**Fig. 3E**). In summary, HBV neutralization requires VIR-3434 as a full-length IgG and is likely mediated via sequestration of virions into SVP-containing immune complexes.

### HBC34/VIR-3434 prevents HBV spread and decreases circulating HBsAg *in vivo* in liver-chimeric mice

Based on its potent and broad *in vitro* neutralization capacity, we tested the effectiveness of HBC34/VIR-3434 in an *in vivo* model of HBV infection. As the specific tropism of HBV for human hepatocytes does not allow infection of wild-type mice, we used a well-established human liver-chimeric mouse model, in which PHH are transplanted into the liver of uPA/SCID beige (USG) mice that lack B and T cells [27, 28]. These mice were infected intraperitoneally with HBV (genotype D) and treated with 1 mg/kg HBC34 (the parental mAb of VIR-3434 with human Fc) or 1 mg/kg HBIG twice per week or 1 µg/ml oral in drinking water of NRTI Entecavir (ETV) during the viral spreading phase starting at 3 weeks post-infection (**Fig. 4A**). After 9 weeks of infection (6 weeks of treatment), serum levels of HBV DNA (i.e., viremia) had increased ∼3 logs above baseline (BL) in mice that were not treated or were treated with HBIG or a control mAb. ETV reduced HBV viremia to below detectable levels at 3 and 6 weeks of treatment, while administration of HBC34 limited HBV viremia to below BL (**Figs. 4B** and **S4A**). Serum HBsAg levels increased ∼2 logs above BL during the treatment period in the control and HBIG treated groups (**Figs. 4C** and **S4B**). HBC34 led to a reduction in serum HBsAg levels to ∼1 log below BL at 6 weeks of treatment. Although HBC34 binds to the antigenic loop of HBsAg, concentrations of up to 50 µg/ml of HBC34 did not interfere with the assay used for HBsAg quantification (Abbott Architect System) in this study (**Fig. S4C**). Analysis of livers after 6 weeks of treatment also demonstrated high efficacy of HBC34 to prevent the increase of intrahepatic viral loads (HBV pgRNA, total HBV RNA, and total HBV DNA), as values were comparable to those at the start of treatment (**Fig. 4D**). In contrast, treatment with HBIG at the same dose as HBC34 did not limit the intrahepatic increase of HBV DNA and RNA. HBV core antigen (HBcAg) staining of mouse liver sections from untreated animals showed a dramatic spread of HBV from few infected cells at 3 weeks post-challenge to infection of nearly all liver-resident human hepatocytes after 9 weeks (**Fig. 4E**). While HBIG showed similar levels of HBcAg positive cells compared to untreated controls, HBC34 or ETV treatment limited intrahepatic spread of HBV (**Fig. 4E**). Given the relatively low potency of HBIG in comparison to HBC34/VIR-3434 (∼1,000-12,000x less potent, see **Figs. 1A** and **1C-D**), the lack of efficacy of HBIG in this study was likely due to the sub-effective dose of HBIG administered to the mice [29]. Our results show that HBC34/VIR-3434 is highly efficacious *in vivo* at blocking the infection of new hepatocytes, thereby limiting the increase in viremia and reducing the level of circulating HBsAg.

**FIGURE 4.**
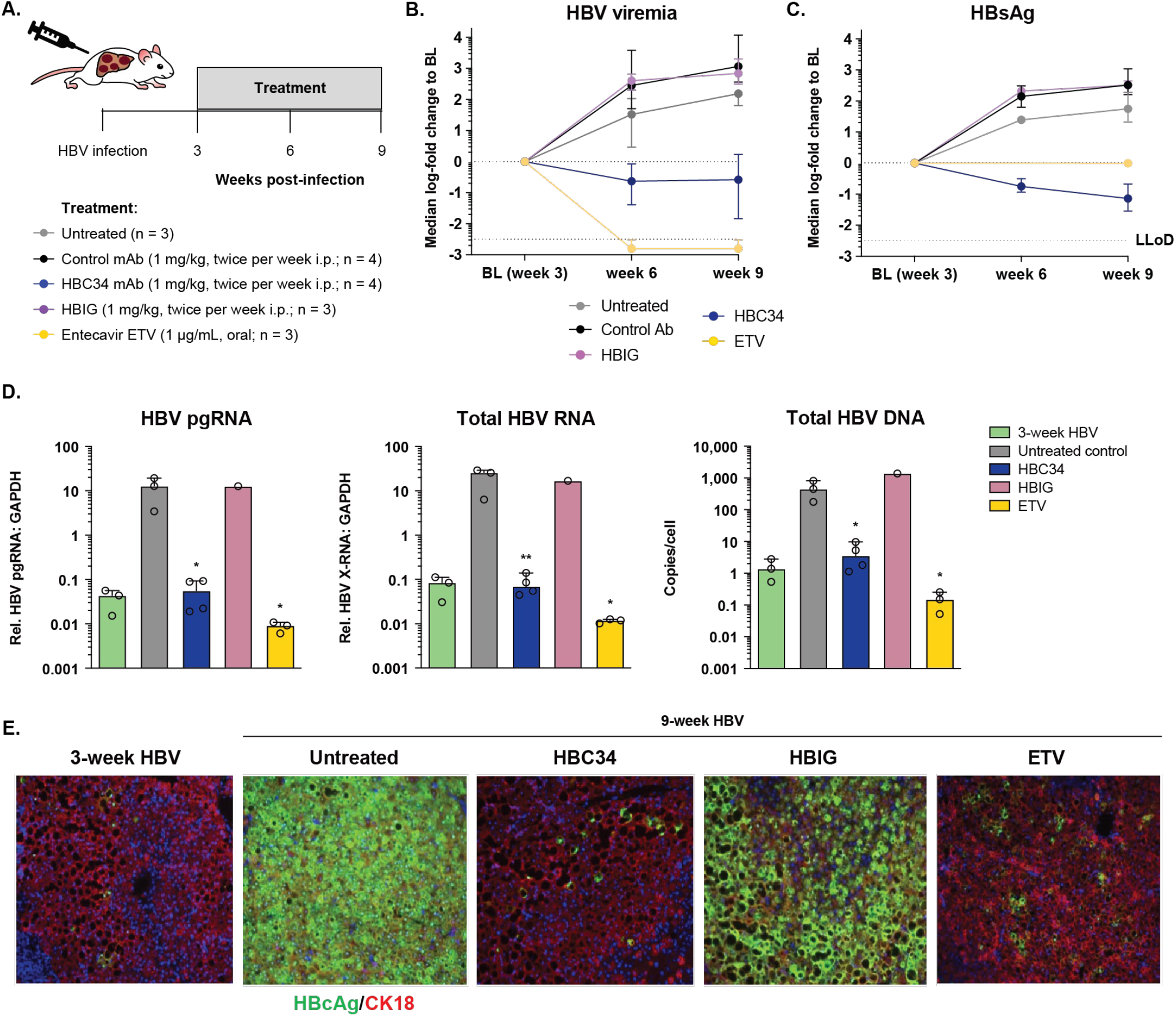
HBC34/VIR-3434 prevents HBV spread and decreases circulating HBsAg in liver-chimeric mice. (A) Human liver-chimeric USG beige mice were infected with HBV (genotype D). Three weeks post infection, at the onset of viral spreading phase, treatment was started twice weekly at 1 mg/kg intraperitoneally with (i) HBC34 (the parental mAb of VIR-3434), (ii) HBIG or (iii) a control mAb or were treated with Entecavir (ETV) at 1 µg/ml in drinking water. Treatment was continued until week 9 post HBV inoculation, when viral infection was spread throughout the human hepatocytes. (B) HBV viremia (HBV DNA) and (C) HBsAg were assessed in serum by qPCR and ELISA, respectively. The mice were sacrificed 9 weeks post infection and intrahepatic HBV pgRNA, total HBV RNA (HBx region), and total HBV DNA (D) were assessed by (RT-)qPCR. Liver sections were immunostained (E) for HBcAg and CK18 as marker for human hepatocytes. In (D), each circle represents one animal. Shown is the median ± range. Statistical differences relative to the untreated control were analyzed by one-way ANOVA. p-value * p ≤ 0.05, ** p ≤ 0.01.

### HBC34/VIR-3434 reduces HBV and HDV viremia in chronically infected human liver-chimeric mice

We next tested the ability of HBC34/VIR-3434 to reduce HBV infection in human liver-chimeric mice with an established infection with high HBV viremia and antigenemia. Mice were infected with HBV for 8 to 12 weeks (see **Fig. 5**) and either not treated or treated with HBC34 (1 mg/kg), with the approved polymerase inhibitor Lamivudine (LAM) at 0.4 mg/ml in drinking water, or with a combination of HBC34 and LAM (**Fig. 5A**). Both drugs alone reduced HBV DNA viremia (∼1 log) after 6 weeks of treatment (**Figs. 5B** and **S5A**). The combination of HBC34 and LAM showed enhanced effectiveness in decreasing viremia compared to the individual agents (greater than 2 logs after 2 weeks treatment and at all subsequent timepoints). Treatment with HBC34 alone achieved a significant reduction of serum HBsAg levels (nearly 2 logs below BL after 6 weeks of treatment); as expected, serum HBsAg levels in LAM treated mice remained close to BL levels throughout the course of treatment (**Figs. 5C** and **S5B**). Notably, the two HBC34 treated animals with lower HBsAg BL levels showed higher reduction and achieved undetectable levels of HBsAg (**Fig. S5B**).

**FIGURE 5.**
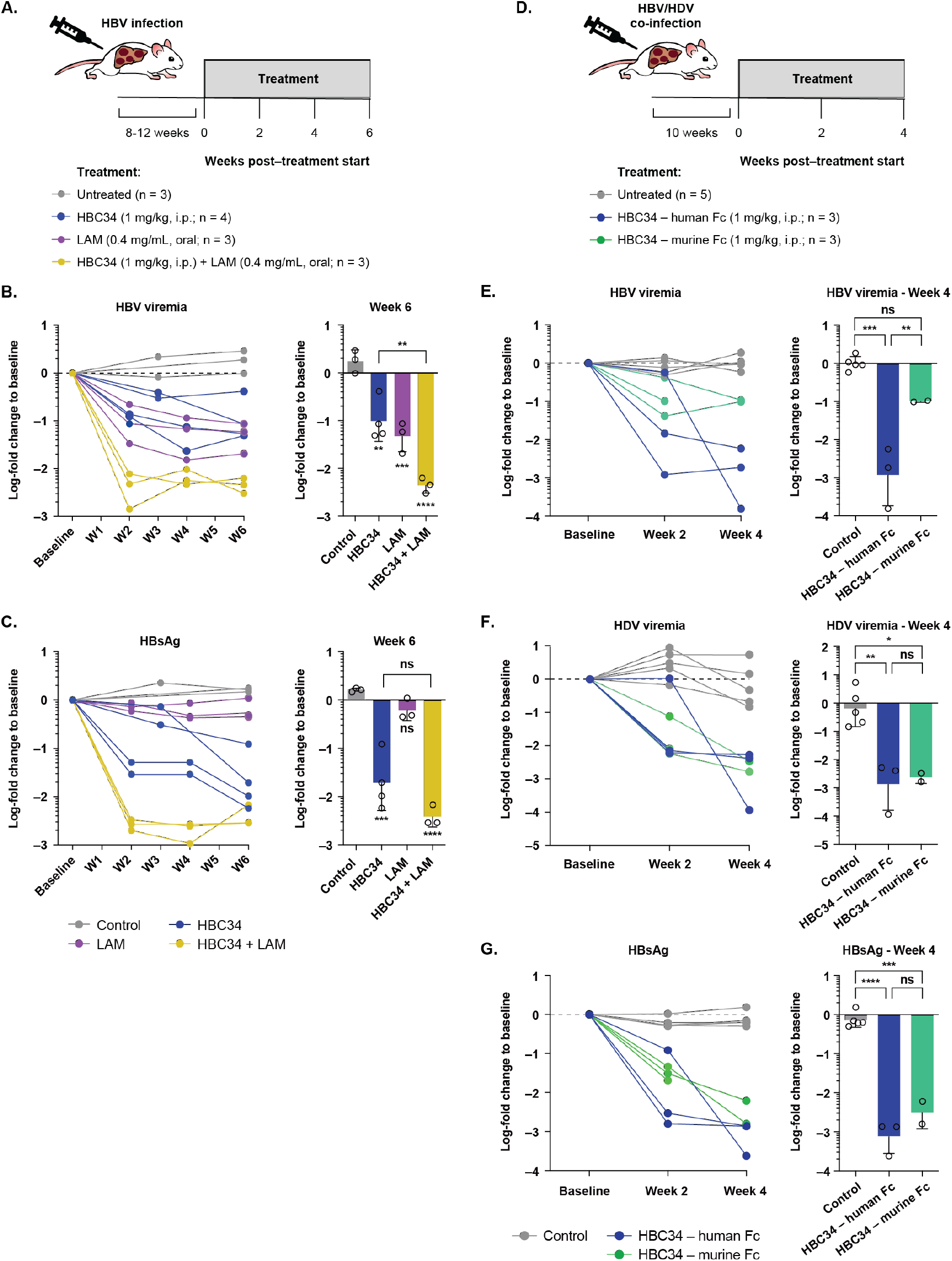
HBC34/VIR-3434 reduces HBV and HDV viremia as well as circulating HBsAg in vivo in chronically infected liver-chimeric mice. (A) Human liver-chimeric USG mice were infected with HBV (genotype D) for 8-12 weeks until stable infection levels were achieved. Mice were treated for 6 weeks twice per week with 1 mg/kg HBC34 (the parental molecule of VIR-3434) intraperitoneally, 0.4 mg/ml lamivudine in drinking water or with both drugs in combination. (B) HBV viremia and (C) HBsAg were assessed in serum by qPCR and ELISA. Graph shows results from 2 independent experiments. (D) Human liver-chimeric mice were co-infected with HBV (genotype D) and HDV (genotype 1) for 10 weeks until stable co-infection was achieved. Mice were treated for 4 weeks with HBC34 either carrying the native human or an engineered murine Fc portion. (E/F) HBV/HDV viremia and (G) HBsAg were assessed in serum by qPCR and ELISA. One animal in the murine Fc group was sacrificed at week 2. Each circle represents one animal. Shown is the mean ± SD. Statistical differences were analyzed by one-way ANOVA. p-value * p ≤ 0.05, ** p ≤ 0.01, *** p ≤ 0.001, **** p ≤ 0.0001, ns p > 0.05.

We then investigated the efficacy of HBC34 in the context of HBV/HDV co-infection. Human liver-chimeric mice that had been stably co-infected with both HBV and HDV (10 weeks after challenge) were treated with HBC34 (1 mg/kg, intraperitoneally twice per week) either carrying wild-type constant regions of the original human IgG1 or a murine IgG2a, which matches the murine FcγRs expressed in this model (**Fig. 5D**). Importantly, human or murine HBC34 reduced both HBV (2.7 log or 1 log reduction) and HDV (2.4 log or 2.6 log reduction) viremia within 2-4 weeks of treatment (**Figs. 5 E-F** and **S5**). In addition, both mAb variants significantly reduced serum HBsAg levels (2.9 log or 2.5 log reduction) (**Figs. 5G** and **S5**). Intrahepatic total HBV DNA, HBV RNA transcripts, and HDV RNA did not change substantially in HBC34-treated mice, indicating that the mAb treatment did not reduce the number of infected cells in this model (**Fig. S5G**). Taken together, our data demonstrate that HBC34/VIR-3434 is highly effective in reducing viremia and circulating HBsAg in mouse models of chronic HBV infection and HBV/HDV co-infection.

## Discussion

In acute viral infections, the use of mAbs is well-established in prophylaxis against respiratory syncytial virus infection of neonates or in post-exposure prophylaxis against Rabies virus infection [30]. For early therapy, potent neutralizing mAbs were recently approved against Ebola virus infections [31], and the Coronavirus Disease 2019 (Covid-19) pandemic showcased that mAbs provide clear clinical benefits [32]. For therapy targeting chronic viral infections, several ultrapotent, broadly neutralizing mAbs are in clinical development against human immunodeficiency virus 1 [30]. In this study, we set out to discover a broadly neutralizing mAb for treatment of CHB. We identified the human mAb VIR-3434 binding with high affinity to HBsAg and neutralizing HBV with broad potency. VIR-3434 had a >12,000-fold higher potency than HBIGs in vitro, which are well-established as post-exposure prophylaxis for newborns to HBV+ mothers [33] as well as in preventing re-infection following liver transplantation [34]. In exploratory clinical studies of patients with CHB, a monthly high dose of HBIG reduced HBsAg levels and induced seroconversion to anti-HBs positive after one year in few individuals with low BL HBsAg (<500 IU/ml) [29, 35]. Monoclonal anti-HBs antibody therapies have been previously evaluated and found to be safe in early, small clinical trials. A human anti-HBs AGL mAb, GC1102, effectively lowered HBsAg by 2 to 3 log_10_ IU/mL and was well tolerated in a Phase 1 study in patients with CHB, with no evidence of serious sequelae, such as immune complex disease [36]. A mixture of two human anti-HBs AGL mAbs, HBV-AB^XTL^ (HepeX-B), significantly reduced HBsAg and HBV-DNA and had a favorable safety and tolerability profile at up to 4 doses of 80 mg administered weekly, with no signs of immune complex disease or hepatotoxicity in 27 patients with CHB [37]. Thus, anti-HBs mAbs may provide promising treatment options for patients with CHB.

We found that VIR-3434 binds to a conformational epitope within the antigenic loop of HBsAg, which forms dimers that protrude from the surface of SVPs and virions [38]. The epitope consists of two highly conserved non-overlapping regions. VIR-3434 binding and neutralization was confirmed for all HBV genotypes (A-H) and a variety of naturally occurring HBsAg variants. In this study, reduced in vitro activity was observed for 2 of 19 HBsAg variants tested (T123N/C124R and G145R). The T123N/C124R double-mutation was associated with a reduced viral fitness as reflected by a reduced HDV titer in vitro, consistent with published results [26]. G145R is a well-described vaccine-escape mutation, which has been observed in infants born to HBV+ mothers who received the HBV vaccine (with or without HBIGs) [39]. The general prevalence of G145R is low (Suppl. Fig. S2) and correlates with specific genotypes C and G. However, published studies selected G145R upon treatment of HBV-infected liver-chimeric mice with HBsAg-specific mAbs [40]. Of note, we did not observe viral rebound in any of the liver-chimeric mice treated with HBC34/VIR-3434 throughout the course of the study. The emergence of mAb-selected HBV escape mutants in patients with CHB may be mitigated by combination treatment of mAb with additional therapeutic agents (e.g. NRTI) to inhibit ongoing replication during the course of treatment.

HBV enters the hepatocyte in a two-step process: initial reversible attachment to HSPGs via the preS1 and AGL regions of HBsAg [5, 41], then the myristoylated preS1 region interacts with the NTCP receptor with high affinity [6]. VIR-3434 neither blocked the interaction of HBsAg with HSPGs nor inhibited viral attachment, indicating no interference with the preS1 region binding to NTCP. Yet the neutralizing activity of VIR-3434 is similar to Ma18/7, currently one of the most potent preS1-specific neutralizing mAbs [42]. Compared to the preS1 epitope of Ma18/7 only present on L-HBsAg of virions and SVP filaments, the epitope of VIR-3434 has higher abundance in vivo, as it is present on the AGL in L-, M- and S-HBsAg on virions, filaments, and spherical SVPs. This comparison of neutralization potency and epitope abundance suggests either a much higher affinity of VIR-3434 or a mode of neutralization that does not require blocking every epitope on HBV virions. We further observed that neutralization by VIR-3434 requires the full-length IgG. In contrast, the Fab demonstrated no neutralizing activity despite binding HBsAg with high affinity. Thus, we hypothesize that the cross-linking of virions and SVPs into immune complexes, as we observed in TEM imaging, is likely required for VIR-3434-mediated neutralization.

In addition to neutralizing viral entry, the formation of immune complexes leads to the clustering of mAb Fcs that is required for high avidity interaction with low-affinity FcgRs. Fc-mediated effector functions may explain the HBC34/VIR-3434-mediated rapid reduction of HBsAg (up to 2 logs) and HBV viremia (1 log) within 2-6 weeks of treatment of chronically HBV-infected human liver-chimeric USG mice. VIR-3434 may opsonize HBsAg SVPs and infectious virions and target them to FcgR-expressing phagocytes, such as monocytes, macrophages, dendritic cells or neutrophils. In a prior study, an anti-HBs mAb (CRL-8017) accelerated the uptake of HBsAg into primary human monocytes, classical dendritic cells, neutrophils, and B cells in vitro [43]. Similarly, treatment with an anti-HBs murine mAb (E6F6) led to rapid clearance of serum HBsAg (>2 log) and HBV DNA via Kupffer cells, macrophages, and neutrophils in several mouse models of HBV infection [44]. As interactions with FcgRs mediated the clearance of HBsAg and virions [44], effector functions are likely pivotal for mAb-mediated therapy of CHB.

To extend serum half-life, the Fc region of VIR-3434 was engineered to include the LS mutation to increase IgG1 binding to human FcRn in endosomes and thus increase recycling into the serum [16-18]. In addition, the Fc region of VIR-3434 was engineered using the GAALIE/XX2 mutation to increase binding to human activating FcγRIIa and IIIa, while decreasing binding to the inhibitory FcγRIIb [19]. This engineered Fc is designed to boost phagocytosis of HBsAg and HBV virions and augment antigen presentation. Previously, the GAALIE/XX2-modified Fc of an anti-HA stem antibody increased the prophylactic potency against influenza A virus infection compared to the mAb carrying a human wild-type Fc in mice transgenic for human FcγRs and FcRn [20]. This increased protection was mediated via fast priming of naïve CD8+ T cells, providing evidence for a mAb-mediated “vaccinal effect”. In human transgenic FcγR and FcRn mice, the GAALIE Fc increased protection in therapeutic models of mAbs against SARS-CoV-2 virus infection [21] or cancer metastasis in the lung [19]. While the tested wild-type human and mouse Fcs of HBC34/VIR-3434 both can interact with the murine FcgRs and FcRn, the GAALIE mutation specifically increases binding only to human FcgRs. The USG mouse model lacking T and B cells and carrying myeloid effector cells with mouse FcgRs thus is not suited to address the benefits of the GAALIE Fc. Future studies using primary human immune cells in vitro, mouse models transgenic for human FcgRs and FcRn, and clinical trials will need to determine whether VIR-3434 can induce a vaccinal effect and mediate long-term anti-HBV immunity.

Beyond HBV monoinfection, more than 12 million chronic HBV carriers are co-infected with HDV worldwide and have an even higher risk of developing liver cirrhosis and hepatocellular carcinoma. Current treatment options are limited to PEG-IFNα, which is associated with only a low frequency of sustained responses, and daily injection of the viral entry inhibitor bulevirtide that has gained conditional marketing approval in the EU [11]. Here, we show that VIR-3434 neutralizes HDV infection in vitro and efficiently reduces HDV viremia in human liver-chimeric USG mice. Entry inhibition and removal of HDV virions/HBsAg from circulation support the potential of VIR-3434 to be an efficacious treatment for patients chronically co-infected with HBV and HDV.

In conclusion, VIR-3434 is a highly potent, half-life extended and effector function-enhanced mAb that binds the antigenic loop present in all forms of HBsAg and neutralizes HBV as well as HDV across all genotypes. VIR-3434 integrates three potential modes of action: (a) inhibition of HBV and HDV entry into new hepatocytes, (b) reduction of circulating HBsAg, and (c) delivering HBsAg to antigen-presenting cells that could reinvigorate adaptive T and B cell responses and long-term immunity via a vaccinal effect. Consequently, the half-life extended VIR-3434 provides a promising therapeutic option for the treatment of patients chronically infected with HBV or also co-infected with HDV and has the potential to induce a functional cure.

## Abbreviations

Anti-HBs: Antibody directed against HBsAg
AGL: Antigenic loop
BL: Base line
BLI: Bio-layer interferometry
CHB: Chronic hepatitis B
ETV: Entecavir
FcRn: Neonatal Fc receptor
FcγRs: Fc gamma receptors
HBIG: Hepatitis B immunoglobulins
HBsAg: Hepatitis B surface antigen
HBV: Hepatitis B virus
HDV: Hepatitis delta virus
HSPG: Heparan sulfate proteoglycan
mAb: Monoclonal antibody
NRTIs: Nucleos(t)ide reverse transcriptase inhibitors
NTCP: Sodium taurocholate co-transporting polypeptide
PEG-IFNa: Pegylated-interferon alpha
PHH: Primary human hepatocytes
SD: Standard deviation
SVP: Subviral particle
TEM: Transmission electron microscopy
USG: uPA/SCID beige

## Acknowledgements

MD, ML and SU received funding from the German Center for Infection Research (BMBF-DZIF: TTU-Hepatitis 05.820; 05.822: 05.714). The German Center of Infection Research (DZIF) TTU Hepatitis provided the infrastructure “Professorship Translational Virology” to SU.

## Author contributions

Conceived study: F.A.L., E.C., F.B., M.D., D.C., M.A.S. Isolated and characterized mAbs: E.C., F.B., F.Z., S.B., G.L., S.J., Performed in vitro virological experiments: F.A.L., J.Z., H.K., H.I., A.S. Performed in vivo experiments and virological measurements: T.V., M.L. SPR/EM experiments: L.E.R, D.M.B. Bioinformatic analysis: L.B.S., A.T. Manuscript writing: F.A.L, T.V., F.B., J.N., M.D., D.C., M.A.S. Supervision and manuscript editing: N.P., S.U., A.T., A.L.C., G.S., L.A.P., C.M.H., M.D., D.C. and M.A.S.

## Data availability statement

All source data that support the findings of this study are available from the corresponding authors upon reasonable request.

## Supplemental Material Online

**Detailed Methods** can be found online.

**Supplemental Figures S1-S5** can be found online.

**Supplemental Table S1** can be found online.

