## Supplemental Methods and Data for "The potent broadly neutralizing antibody VIR-3434 controls Hepatitis B and D Virus infection and reduces HBsAg in humanized mice"

### Supplementary Information

Florian A. Lempp, Tassilo Volz, Elisabetta Cameroni, Fabio Benigni, Jiayi Zhou, Laura E. Rosen, Julia Noack, Fabrizia Zatta, Hannah Kaiser, Siro Bianchi, Gloria Lombardo, Stefano Jaconi, Hasan Imam, Leah B. Soriaga, Nadia Passini, David M. Belnap, Andreas Schulze, Marc Lütgehetmann, Amalio Telenti, Andrea L. Cathcart, Gyorgy Snell, Lisa A. Purcell, Christy M. Hebner, Stephan Urban, Maura Dandri, Davide Corti, and Michael A. Schmid

### Supplementary Materials and Methods

#### HBV production

HBV was derived from HepAD38 cells [1], which harbor a tetracycline-inducible HBV more-than-genome-length genomic integrate (genotype D). Cell culture supernatants were collected for virus purification beginning at day 14 after removal of tetracycline. Virus in the supernatant was concentrated 50-fold by precipitating viral particles with 6% PEG-8000 (Sigma-Aldrich) overnight at 4°C. Precipitates were recovered by centrifugation and resuspended in 20 mM Tris/140 mM NaCl buffer containing 10% FBS. Virus stock was aliquoted and stored at -80°C.

#### HDV production

Huh7 cells were seeded in 10-cm dishes at 5x10$^{6}$ cells per dish. The next day, cells were co-transfected with the plasmid pcDNA-HDVgt1Ethiopia (encoding for the HDV antigenome [2]) and a plasmid encoding a subgenomic fragment of HBV (HB2.7) encompassing all HBV surface antigen ORFs [3]). HB2.7 constructs of different HBV genotypes were previously described, [2] except for genotype A (EU054331), which was newly synthesized. HB2.7 constructs bearing single amino acid variants were all cloned in the backbone of HBV genotype D (NC_003977). Medium was changed one day post transfection and every second or third day thereafter until day 10. The HDV-containing cell supernatant was collected at day 12 post transfection, centrifuged at 2,000 rpm for 5 min to remove cellular debris, aliquoted and frozen at -80°C.

#### HBsAg binding ELISA

Half-area 96-well plates were coated with 25 µl/well HBsAg adw (Prospec, hbs-872-b), HBsAg adr (Prospec, hbs-875-b) or HBsAg ayw (Meridian, R86870) at 1 µg/ml and incubated overnight at 4°C. Plates were washed three times with PBS-T using an automated washer. Blocking solution (PBS with 1% BSA) was added and plates were further incubated for 1 hour at room temperature (RT). Blocking solution was removed and plates were washed again. Then 25 µl/well of serial 1:3 mAb dilutions (10 µg/ml to 0.5 ng/ml) in blocking buffer were dispensed and plates were incubated 90 min at RT. Plates were then washed 4 times with PBS-T (220 µl/well). The HRP-conjugated secondary antibody reagent goat anti-human IgG (Jackson Immuno, 109-035-098) was added to each well at 0.16 µg/ml in blocking buffer and further incubated for 45 min at RT. After 4 washes with PBS-T, 40 µl/well of Sureblue (TMB) ELISA substrate solution was dispensed in each well and plates were developed for 15 min at RT. The reaction was stopped with 1% HCl, and the OD was read at 450 nm in an ELISA reader (Bio-Tek, ELx808).

#### ELISA competition assay

Selected antibodies were labeled with biotin and tested by ELISA in a matrix competition assay, in which unlabeled antibodies were incubated first on HBsAg-coated plates, followed by the addition of a limiting concentration of biotinylated antibodies. Binding was revealed with alkaline phosphatase-conjugated streptavidin. If two epitopes overlap, or the areas covered by the arms of the two antibodies overlap, competition should be almost complete. Weak inhibitory or enhancing effects may reflect a decrease in affinity owing to steric or allosteric hindrance.

**Flow cytometry based HBsAg binding**

The HBsAg protein coding sequences of different genotypes were derived from the public domain database GenBank NCBI and cloned into cell expression vectors (phCMV1) under the control of the human CMV promoter (genotypes A (AM282986), B (D23678), C (AB117758), D (AB126581), E (AB205192), F (X69798), G (AF160501), H (AY090454), I (AF241409), J (AB486012), all single variant substitutions were made in the genotype D (AB126581) backbone). Expi293 cells were seeded in 10 ml medium in 125 ml Flask at $2.5x10^6$ cells/ml on the day of transfection. HBsAg-encoding plasmids were diluted in 0.5 ml Opti-MEM-I and mixed with an equal volume transfection reagent solution (3 µM PEI MAX 40 in Opti-MEM-I) and complexed for 20 min at RT. The transfection mix was added dropwise to cells with gentle swirling followed by a 48-hour incubation at 37°C, 5% CO2. Two days post-transfection, cells were harvested by 5 min centrifugation at 400 rcf at room temperature, the pellet was dislodged, and cells were washed twice with PBS. Cells were resuspended in 20 ml fixative solution incubated 20 minutes on ice and washed again twice with PBS. The pellets were then resuspended in permeabilization solution to a final concentration of $0.5x10^6$ cells/ml and 100 µl of this suspension were dispensed in each well of a 96-well round bottom plate. After 20 min incubation on ice dilutions of the mAbs were added to each well (50 µl/well). Dilution series were prepared separately in permeabilization solution by performing 3-fold dilution series from a starting concentration of 10 µg/ml. After 30 min incubation on ice the cells were washed twice with permeabilization buffer (150 µl/well) and 50 µl of Alexa Fluor 647-labelled secondary antibody (diluted to 2.5 µg/ml in permeabilization solution) were added to the cells and incubated for 30 min on ice. Cells were washed two more times with permeabilization solution and finally resuspended in 150 µl permeabilization solution. Mean fluorescence intensity signal was quantified with a cytofluorimeter (Beckton Dickinson). The mean fluorescence intensity was determined after gating on the transfected cell populations as determined by comparison with the signal obtained in the mock controls. Flow cytometry data analysis was performed using Flowjo software. Data was plotted using GraphPad Prism 8.0.

**HBV neutralization (HepaRG)**

HepaRG cells were grown in Williams' medium E supplemented with 10% heat-inactivated fetal calf serum, 50 U of penicillin/ml, 50 ug of streptomycin/ml, 5 ug of insulin/ml, and 50 uM hydrocortisone hemisuccinate. Cells were passaged every 2 weeks at a ratio of 1:5. Fourteen days before infection, cell differentiation was induced by adding 1.5% dimethyl sulfoxide to the maintenance medium. The medium was exchanged every 3 days. For infection, differentiated HepaRG cells ($1x10^6$ cells/well of a 12-well plate) were incubated with a 20-fold dilution of the concentrated virus stock that had been preincubated with antibody dilutions. Virus-antibody mixtures were added to the cells in medium containing 4% polyethylene glycol 8,000 for 16 h at 37°C. At the end of the incubation, the cells were washed three times and further cultivated.

Medium was changed every 3 days. To quantify the infection, amounts of hepatitis B surface antigen (HBsAg) and hepatitis B e antigen (HBeAg) secreted into the culture supernatant from day 7 to 11 postinfection (p.i.) were determined by enzyme-linked immunosorbent assay ([ELISA] AxSYM; Abbott).

**HBV neutralization (PHH)**

Primary human hepatocytes (PHH, Thermo Fisher Scientific) were seeded at 58,000 cells/well in collagen-coated 96-well plates according to the manufacturer's instructions. Five hours post seeding, at ~90% confluence, infection was initiated. Infection mix was prepared by diluting the concentrated HBV stock virus at 1:30 in PHH maintenance medium (Williams E, primary hepatocyte maintenance supplements [Thermo Fisher Scientific], 2% FBS, 2% DMSO). Five-fold, eight-step serial dilutions of the antibodies were prepared in PBS starting at 62.5 μg/mL (VIR-3434, Ma18/7) or 100 mg/mL (HBIG [Sigma-Aldrich])) and diluted 1:10 into the infection mix. Infection control wells contained infection mix with PBS alone. Virus-antibody complexes were allowed to form by 30-minute incubation at 37°C, then 40% PEG-8000 was added to a final PEG concentration of 4% and mixed well. One hundred microliters from each well of the virus-antibody complex plates were pipetted in triplicates onto the cells. One day post infection, the inoculum was removed, cells were washed with PBS and fresh PHH maintenance medium was added. Medium was exchanged every 2-3 days post infection. At day 7 post infection, viral markers in the cell supernatant were quantified by HBeAg/HBsAg CLIA according to the manufacturer's instructions (Autobio Diagnostics).

**HBV neutralization competition with exogenous HBsAg**

PHH were seeded at 58,000 cells/well in collagen-coated 96-well plates according to the manufacturer's instructions. Five hours post seeding, at ~90% confluence, infection was initiated. Infection mix was prepared by diluting the concentrated HBV stock virus 1:30 in PHH maintenance medium. Five-fold, five step serial dilutions of the exogenous HBsAg preparations (Yeast HBsAg [Prospec, HBS-872], PLC HBsAg [cell supernatant of PLC/PRF/5 cells, ATCC CRL-8024]) were prepared in PBS starting at 16,662 IU/mL and diluted 1:8.33 into the infection mix. Five-fold, seven step serial dilutions of VIR-3434 were prepared in PBS starting at 12.5 μg/mL and diluted 1:10 into the HBsAg/infection mix dilutions in a checkerboard format. Infection control wells contained only infection mix with neither exogenous HBsAg nor VIR-3434. Virus/HBsAg-antibody complexes were allowed to form via 30 minutes incubation at 37°C, then 40% PEG-8000 was added to a final PEG concentration of 4% and mixed well. One hundred microliters from each well of the virus-antibody complex plates were pipetted onto the cells in duplicate plates. One day post infection, the inoculum was removed, cells were washed with PBS and fresh PHH maintenance medium was added. Medium was exchanged every 2-3 days post infection. At day 8 post infection, viral markers in the cell supernatant were quantified by HBeAg/HBsAg CLIA according to the manufacturer's instructions (Autobio Diagnostics).

### HDV neutralization

Huh7-NTCP cells were seeded in transparent, clear-bottom 96-well plates at 20,000 cells/well in 100 µL medium and cultured overnight at 37°C to reach ~90% confluency. Infection mix was prepared by diluting the respective HDV virus stock in cDMEM containing 2% DMSO. When comparing neutralization of HDV enveloped with HBsAg of different HBV genotypes, virus preparations were normalized across genotypes based on their HBsAg levels determined by ELISA. Serial dilutions of mAbs were prepared in PBS. Virus-antibody complexes were allowed to form by 30-minute incubation at 37°C, then 40% PEG-8000 (Sigma-Aldrich) was added to a final PEG concentration of 4% and mixed well. One hundred microliters from each well of the virus-antibody complex plates were pipetted in triplicates onto the cells. After 16 hours incubation, the infectious inoculum was removed, cells were washed twice with PBS and 100 µL medium (cDMEM with 2% DMSO) added to each well. Medium was changed again at days 4 and 5 post infection. At day 7 post infection, viral infection was quantified by immunofluorescence staining for HDAg. Cell supernatant was removed, cells were washed once with PBS and fixed with 50 µL/well 4% PFA for 30 min at RT followed by two PBS washes and permeabilization with 50 µL/well PBS/0.25% Triton-X100 for 30 min at RT. Staining was performed with 50 µL/well mouse-anti-HDAg (clone FD3A7, kind gift from Stephan Urban) primary antibody diluted 1:2000 in PBS with 5% milk powder for 3 h at RT. After primary antibody staining, cells were washed three times with PBS (5 min incubation time each) and incubated with 50 µL/well goat-anti-mouse-Alexa647 diluted 1:2000 in PBS/5% milk powder with 2 µg/mL Hoechst dye for 1h at RT. Cells were washed three times with PBS and imaged on a Cytation 5 automated fluorescence microscope (Biotek) acquiring 12 images per well with the 4x objective in two channels using DAPI and Cy5 filter sets. HDAg-positive cells were counted using the manufacturer's software.

### HBsAg Western Blot

Samples were mixed with 4x Laemmli Buffer (Bio-Rad) containing β-Mercaptoethanol (Bio-Rad) and then heated at 95°C for 10 minutes. 4-20% precast protein gel (Bio-Rad) was used for electrophoresis and then transferred to a PVDF membrane (Bio-Rad) using semi-dry transfer method (Trans-Blot Turbo Transfer System, Bio-Rad) according to the manufacturer's instructions. The membrane was blocked with 5% BSA in 1X TBST for 1 hour, followed by overnight incubation with the primary antibody at 1:1000 dilution (human-anti-HBs (clone HBD87), rabbit-anti-preS1 (H849)). After washing with 1X TBST, the membrane was incubated with HRP conjugated secondary antibody (1:5000) for 1 hour. After washing, the membrane was incubated in ECL reagent (Bio-Rad) for 1-2 minutes. The signals were detected using a chemiluminescence imaging system (Azure Biosystems).

### Epitope mapping

Epitope mapping was performed by Pepscan Presto BV (Lelystad, The Netherlands). Epitope was identified by using a library of 650 linear and looped peptides designed to cover the entire antigenic loop region of HBsAg (CLIPS Discontinuous Epitope Mapping technology [4]). Target-derived peptides were chemically synthesized and immobilized into defined 3D structures. The binding of antibody to each of the synthesized peptides was tested in a pepscan-based ELISA. The peptide arrays were incubated with primary antibody solution (overnight at 4°C). After washing, the peptide arrays were incubated with a 1/1000 dilution of an appropriate antibody peroxidase conjugate for one hour at 25°C. After washing, the peroxidase substrate 2,2'-azino-di-3-ethylbenzthiazoline sulfonate (ABTS) and 20 µl/ml of 3 percent H2O2 were added. After one hour, the color development was measured. The color development was quantified with a charge coupled device (CCD) camera and an image processing system.

**HBV inhibition of binding assay**

HepG2-NTCP cells were seeded in clear-bottom, 24-well plates at 300,000 cells/well and cultured overnight at 37°C to reach ~90% confluency. Infection mix was prepared by mixing respective antibodies with HBV in DMEM containing 2% DMSO. Virus-antibody complexes were allowed to form by 30-minute incubation at 37°C. Cells in the Myrcludex B (MyrB, Genscript) control wells were preincubated with 1 µM MyrB in DMEM for 30 min and 1 µM MyrB was additionally added to the infection mix. 250 µL from each virus-antibody complex was pipetted in triplicates onto the cells. After three hours incubation, the infectious inoculum was removed, and cells were washed three times with PBS with two to three minutes incubation between each wash. Total DNA was isolated from the cells using NucleoSpin DNA RapidLyse (Macherey-Nagel) as described in the manufacturer's protocol using proteinase K digestion. DNA concentration was measured by Nanodrop and adjusted with water according to the lowest concentration. Four microliters of each sample were used in the qPCR reaction with 16 µL of a mastermix containing 3.6 µL of water, 2 µL of primer (5 µM each, for: GTGTCTGCGGCGTTTTATCA, rev: GACAMACGGGCAACATACCT), 0.4 µL of probe oligonucleotide (10 µM, seq: CCTCTKCATCCTGCTGCTATGCCTCATC) and 10 µL of the qPCR NEB Luna Probe 2x reaction mix. qPCR reactions were run and acquired on a QuantStudio 3 Real-Time PCR System (Applied Biosystems). The two-step thermal cycle was programmed for 10 minutes at 95°C as initial denaturation, followed by 40 cycles of 15 sec at 95°C for denaturation, 60 sec at 60 °C for annealing and extension. Standard curve reactions were prepared using known amounts of HindIII-linearized pUC-HBV1.3 plasmid.

**Surface plasmon resonance (SPR)**

Measurements were performed via SPR using a Biacore T200 instrument. A CM5 chip covalently immobilized with Mouse antibody capture kit (Cytiva) was used for surface capture of anti-HBs HBD7-muFc (mAb non-competing with VIR-3434) and subsequent HBsAg capture. Running buffer was HBS-N pH 7.4 (Cytiva) supplemented with 0.01% BSA (Sigma A7030); measurements were performed at 25°C. Experiments were performed with a 3-fold dilution series of VIR-3434 IgG or Fab fragment (0.74, 2.2, 6.7, 20, 60 nM). Data were double reference-subtracted and fit using Biacore Evaluation software to a Heterogeneous Ligand binding model (the data were not fit well by a 1:1 binding model, likely because of heterogeneity in the HBsAg antigenic loop which contains the VIR-3434 epitope). Apparent affinities (KD,app) are reported as an upper bound based on the weakest binding reported by the fit. Results are representative of duplicate measurements and are consistent with additional replicates in alternate experimental formats.

### Transmission electron microscopy (TEM)

Yeast-derived HBsAg particles (ProSpec HBS-872, serotype adw, 30,000 IU/ml, concentration determined using the Elecsys HBsAg II quantitative assay, Cobas e801, Roche) and VIR-3434 IgG or Fab fragment were diluted in PBS (Gibco 10010-031) to a final concentration of 1,500 IU/ml HBsAg and 5 μg/ml VIR-3434 IgG or 3.3 μg/ml VIR-3434 Fab fragment (Fab concentration chosen to match the molar concentration of Fab arms present in the 5 ug/ml IgG sample). An apo sample of 1,500 IU/ml HBsAg was also prepared. Samples were incubated in a 37°C water bath for 60-65 min immediately prior to negative staining. A negatively stained specimen of each sample was prepared on ultrathin amorphous carbon supported on a 200-mesh Cu grid (CF200-CU-UL, Electron Microscopy Sciences, Hatfield, Pennsylvania, USA). (The carbon-coated grid was first placed in a lab-made plasma, glow-discharge device for 10-15 seconds to clean the carbon and render it more hydrophilic.) Stain solution was 1% ammonium molybdate. Specimens were prepared using the following steps: application of 3.5 μL sample to the grid for one minute then blot via filter paper; 2x application (1-2 sec) of 20 μL PBS then blot; 1-2x application (1-2 sec) of stain solution then blot; final application of 20 μL stain solution for 15–20 seconds, blot, dry in air or vacuum. Grids were imaged on a ThermoFisher Tecnai 12 transmission electron microscope operated at 120 kV. At least 20 images were recorded at different positions across each grid using a Gatan UltraScan camera; for presentation, one representative image was chosen from each dataset.

### Generation of humanized USG mice

Human liver chimeric urokinase-type plasminogen activator (uPA)/severe combined immunodeficiency(scid)/beige/interleukin-2 receptor gamma chain negative ($IL2R\gamma^{-/-}$) mice (short USG mice) were generated by transplanting one million thawed cryo-preserved human hepatocytes into homozygous USG mice as previously reported [5]. Repopulation rates were estimated by determining human serum albumin (HSA) in mouse sera (ELISA; Bethyl Laboratories, Biomol GmbH, Hamburg, Germany) and human beta-globin DNA in the liver (qRTPCR; Taqman Gene Expression Assay Hs00758889_s1; Applied Biosystems, Carlsbad, CA, United States). Animals displaying high levels of human chimerism (>2 mg/ml HSA in serum) were used for the study. Mice were sacrificed at different timepoints, as indicated in the results, blood was collected and liver specimens were snap-frozen in chilled isopentane and cryo-conserved at -80°C for histological and molecular analyses. Mice were maintained under specific pathogen free conditions in accordance with institutional guidelines under approved protocols. All animal experiments were conducted in accordance with the European Communities Council Directive (86/609/EEC) and were approved by the City of Hamburg, Germany.

### Mouse infections and treatment

Human liver chimeric USG mice were infected either with HBV (mono-infection) or with HBV and HDV (co-infection). Humanized mice received a single peritoneal injection of infectious serum containing HBV genotype D (1x$10^7$ HBV genome equivalents/mouse) or a mixture of HBV and HDV-genotype 1 (both 1x$10^7$ genome equivalents/mouse), which was also passaged through humanized USG mice. All treatment regimens performed during the viral spreading phase (HBIG, ETV, HBC34 or control Ab) were started 3 weeks after viral inoculation and maintained for six weeks. Mice sacrificed 3 weeks post-infection served as intrahepatic baseline control. To assess therapies in the chronic phase of infection, treatments (HBC34, Lam or HBC34+Lam in combination) were started 8-10 weeks post HBV mono-infection or 10 weeks after HBV/HDV coinfection (HBC34 with murinized or humanized Fc part). Mice received 1 mg/kg HBC34, control or HBIG antibody (Hepatect CP) two times weekly i.p. for the indicated duration. Entecavir was given oral in drinking water supplied with 1 µg/ml and lamivudine (Zeffix, GlaxoSmithKline, Brentford, UK) at 0.4 mg/ml. Controls were left untreated.

### In vivo virological measurements and immunostaining

Quantification of HBsAg from mouse sera was performed using the Abbott Architect platform (Abbott, Ireland, Diagnostic Division). Immunohistochemistry on mouse liver sections was performed as described in [6]. Briefly, acetone fixed frozen liver sections and human-specific antibodies recognizing keratin 18 (Santa Cruz Biotechnology, Dallas, TX, USA) were used to visualize human hepatocytes. HBV-infected hepatocytes were identified using an HBcAg-specific antibody (Dako Diagnostika,Glostrup, Denmark). Nuclear staining was achieved with Hoechst 33258 (Invitrogen). Stained sections were analyzed by fluorescence microscopy (Biorevo BZ-9000, Keyence, Osaka, Japan) using the same settings for all groups.

DNA and RNA were extracted from liver specimens using the Master Pure DNA Purification Kit (Epicentre) and the RNeasy RNA Purification Kit (QIAGEN), respectively. Mouse sera were cleaned up with Qiagen MinElute virus spin Kit.

HBV viremia and intrahepatic HBV DNA levels were determined by qRT-PCR using specific primers and probes (HBV DNA: Taqman Gene Expression Assay Pa03453406_s1, Applied Biosystems) under conditions previously described [7]. Intrahepatic viral RNA loads were determined using primers and probes specific for total HBV RNA (targeting the X-region) (Taqman Gene Expression Assay Pa03453406_s1, Applied Biosystems), pregenomic (pg)RNA [8] and values normalized using the mean of human house-keeping genes *GAPDH (*Hs99999905_m1) and RPL30 (Hs00265497_m1).

HDV qRTPCR was performed as described in [7]. HDV viremia and intracellular HDV RNA was determined via reverse transcription and qRT-PCR using the ABI Fast 1-Step Virus Master (Applied Biosystems, Carlsbad, USA) on an ABI Viia7 (Applied Biosystems). In brief, 5µl mouse sera were cleaned up with Qiagen MinElute virus spin Kit and 5 µl RNA was denatured at 95°C for 10 min, immediately cooled down on ice and reverse transcribed at 50°C for 5 min with HDV specific primers and probes. After inactivation of the reverse transcriptase at 95°C for 20 s, amplification was performed under the following conditions: 40 cycles at 95°C for 3 s and 60°C for 30 s 4. The plasmid pBluescript II SK(þ), containing one copy of the HDV genome, was used as a standard for HDV cDNA quantification.

**A.**

| mAb | Competition with biotinylated mAb: 17.1.41 | 19.79.5 | HBC34 | HBD65 | HBD7 | HBC24 | HBC30 | HBD85 |
|---|---|---|---|---|---|---|---|---|
| **17.1.41** | + | +/– | +/– | +/– | – | +/– | +/– | +/– |
| **19.79.5** | – | + | +/– | – | – | + | – | – |
| **HBC34** | – | + | + | – | – | +/– | – | – |
| **HBD65** | – | – | – | + | +/– | – | + | + |
| **HBD7** | – | – | – | + | + | – | + | + |
| **HBC24** | – | + | +/– | – | – | + | – | – |
| **HBC30** | – | – | – | + | – | – | + | + |
| **HBD85** | – | – | – | + | – | + | + | + |

**B.**

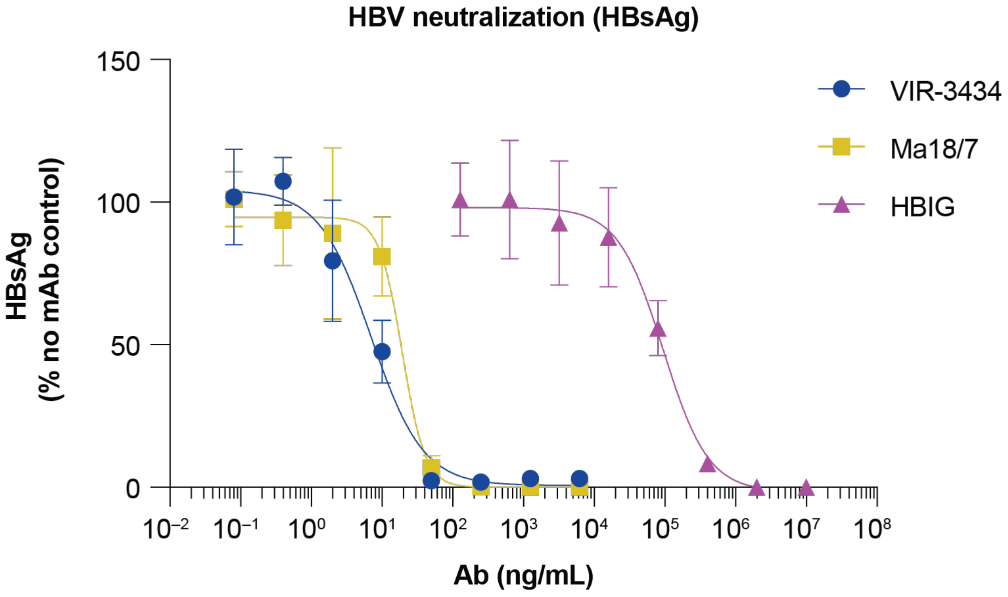

**SUPPLEMENTARY FIGURE S1. Characterization of HBsAg-targeting human monoclonal antibodies.** (A) Competition of mAbs was assessed by ELISA. A filled box (+) indicates that the mAbs compete in both direction of first vs. second binder, (+/–) indicates competition in only one direction, and (–) indicates no competition. (B) HBV neutralization using secreted HBsAg as readout for infection using primary human hepatocytes (PHH).

**A.**

| HBsAg amino acid position | Genotype (number of sequences) | | | | | | | | | |
|---|---|---|---|---|---|---|---|---|---|---|
| | **ALL n=28,330** | **A n=3,474** | **B n=6,361** | **C n=10,348** | **D n=4,948** | **E n=1,018** | **F n=585** | **G n=152** | **H n=129** | **RF n=1,316** |
| 114 | S=84.3, T=15.13 | T=97.11, A=1.3, S=1.1 | S=99.36 | S=98.66 | S=99.35 | S=99.12 | T=98.12 | S=100 | T=96.9, S=3.1 | S=92.93, T=6.77 |
| 115 | T=99.73 | T=99.91 | T=99.67 | T=99.8 | T=99.57 | T=99.8 | T=98.97 | T=100 | T=100 | T=99.77 |
| 116 | T=99.7 | T=99.8 | T=99.86 | T=99.7 | T=99.7 | T=99.9 | T=100 | T=100 | T=100 | T=98.25, A=1.6 |
| 117 | S=99.18 | S=99.83 | S=99.75 | S=98.26, T=1.41 | S=99.82 | S=99.51 | S=99.32 | S=100 | S=100 | S=99.16 |
| 118 | T=95.29, V=2.42, A=1.28 | T=99.68 | T=99.54 | T=97.64, M=1.88 | T=79.45, V=13.62, A=6.56 | T=99.51 | T=100 | T=100 | T=99.22 | T=97.57, V=1.06 |
| 119 | G=99.89 | G=99.74 | G=99.92 | G=99.88 | G=99.92 | G=100 | G=100 | G=100 | G=100 | G=99.92 |
| 120 | P=98.53 | P=98.06, T=1.5 | P=98.03 | P=99.39 | P=97.7, S=1.12 | P=99.31 | P=98.29, Q=1.03 | P=99.34 | P=100 | P=97.72, T=1.29 |
| 121 | C=99.89 | C=99.86 | C=99.95 | C=99.89 | C=99.82 | C=99.9 | C=100 | C=100 | C=100 | C=99.77 |
| 122 | K=70.51, R=29.33 | K=91.38, R=8.45 | K=78.83, R=21.04 | K=98.63, R=1.21 | R=98.09, K=1.7 | R=99.9 | K=97.26, R=2.56 | K=89.47, R=10.53 | K=99.22 | R=49.96, K=49.81 |
| 123 | T=99.35 | T=99.68 | T=99.13 | T=99.38 | T=99.19 | T=99.8 | T=99.83 | T=100 | T=100 | T=99.09 |
| 124 | C=99.81 | C=99.94 | C=99.91 | C=99.77 | C=99.62 | C=100 | C=99.66 | C=100 | C=100 | C=100 |

**B.**

| HBsAg amino acid position | Genotype (number of sequences) | | | | | | | | | |
|---|---|---|---|---|---|---|---|---|---|---|
| | **ALL n=28,329** | **A n=3,474** | **B n=6,361** | **C n=10,348** | **D n=4,948** | **E n=1,018** | **F n=585** | **G n=152** | **H n=129** | **RF n=1,316** |
| 145 | G=98.48 | G=99.19 | G=99.32 | G=97.5, R=1.22, A=1.03 | G=98.54 | G=98.52 | G=99.83 | G=98.03, R=1.97 | G=99.22 | G=99.24 |
| 146 | N=99.85 | N=99.91 | N=99.81 | N=99.82 | N=99.92 | N=99.9 | N=99.83 | N=100 | N=100 | N=99.77 |
| 147 | C=99.84 | C=99.74 | C=99.8 | C=99.86 | C=99.92 | C=100 | C=100 | C=100 | C=100 | C=99.62 |
| 148 | T=99.87 | T=99.94 | T=99.73 | T=99.88 | T=99.96 | T=99.9 | T=99.83 | T=100 | T=100 | T=99.85 |
| 149 | C=99.89 | C=99.88 | C=99.91 | C=99.86 | C=99.9 | C=100 | C=100 | C=100 | C=100 | C=100 |
| 150 | I=99.89 | I=99.88 | I=99.91 | I=99.89 | I=99.9 | I=99.71 | I=100 | I=100 | I=99.22 | I=100 |
| 151 | P=99.92 | P=99.94 | P=99.97 | P=99.91 | P=99.9 | P=99.8 | P=100 | P=100 | P=100 | P=99.7 |
| 152 | I=99.89 | I=100 | I=99.94 | I=99.84 | I=99.86 | I=100 | I=100 | I=100 | I=100 | I=99.85 |
| 153 | P=99.87 | P=99.97 | P=99.92 | P=99.74 | P=99.92 | P=100 | P=100 | P=100 | P=100 | P=100 |
| 154 | S=99.8 | S=99.34 | S=99.94 | S=99.85 | S=99.84 | S=99.9 | S=100 | S=100 | S=100 | S=99.62 |
| 155 | S=99.91 | S=99.94 | S=99.89 | S=99.86 | S=99.96 | S=100 | S=100 | S=100 | S=100 | S=100 |
| 156 | W=99.62 | W=99.86 | W=99.23 | W=99.57 | W=99.94 | W=99.9 | W=99.49 | W=99.34 | W=100 | W=99.85 |
| 157 | A=99.86 | A=99.68 | A=99.86 | A=99.9 | A=99.84 | A=99.9 | A=100 | A=100 | A=100 | A=99.92 |
| 158 | F=97.7, L=2.18 | F=99.86 | F=99.68 | F=99.85 | F=99.57 | F=99.41 | L=99.49 | F=100 | F=99.22 | F=99.85 |

**C.**

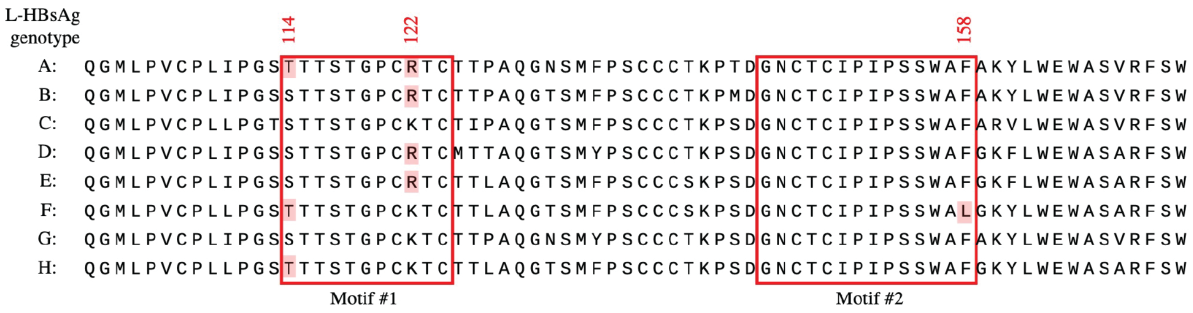

**SUPPLEMENTARY FIGURE S2. Conservation of VIR-3434 epitope.** Percentage of amino acid present per genotype in ≥ 1.0% of sequences in motif #1 (A) and motif #2 (B). Data is based on complete and partial HBV genomes downloaded from HBVdb, n = 28331 (2021 Oct 18). RF: recombinant forms (not assigned to genotypes). (C) Alignment of L-HBsAg antigenic loop sequences from HBV genotypes A-H used in this study. The two amino acid motifs recognized by VIR-3434 are highlighted in red boxes. Amino acid positions 114, 122 and 158 are highlighted and less prevalent amino acids shaded in red.

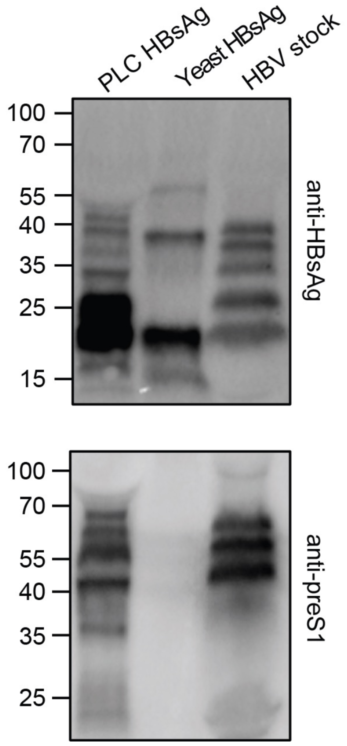

**SUPPLEMENTARY FIGURE S3. Characterization of HBsAg protein preparations in SVPs.** Western Blot analysis probing with an antibody targeting S-HBsAg (top) or preS1 (bottom). HBV virus stock produced in HepAD38 cells was used as positive control.

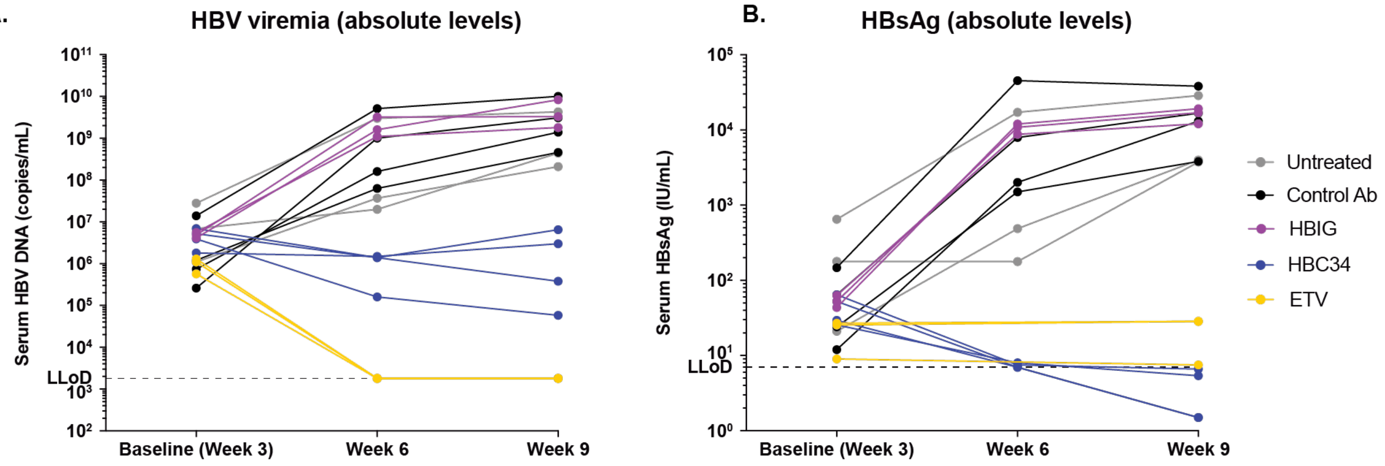

C.

| Mouse serum with HBsAg | Spike-in HBC34 | Abbott Architect measurement |
|---|---|---|
| 6 IU/mL | 0 µg/mL | 5.85 IU/mL |
| 6 IU/mL | 5 µg/mL | 5.79 IU/mL |
| 6 IU/mL | 50 µg/mL | 5.95 IU/mL |

**SUPPLEMENTARY FIGURE S4. Inhibition of viral spread in liver-chimeric mice.** Absolute levels of HBV viremia (A) and HBsAg (B) in all mice over the treatment period. (C) Possible HBC34 interference with HBsAg quantification assay was assessed by spike-in of HBC34 in mouse serum containing 6 IU/ml HBsAg and HBsAg measurement on the Abbott Architect system.

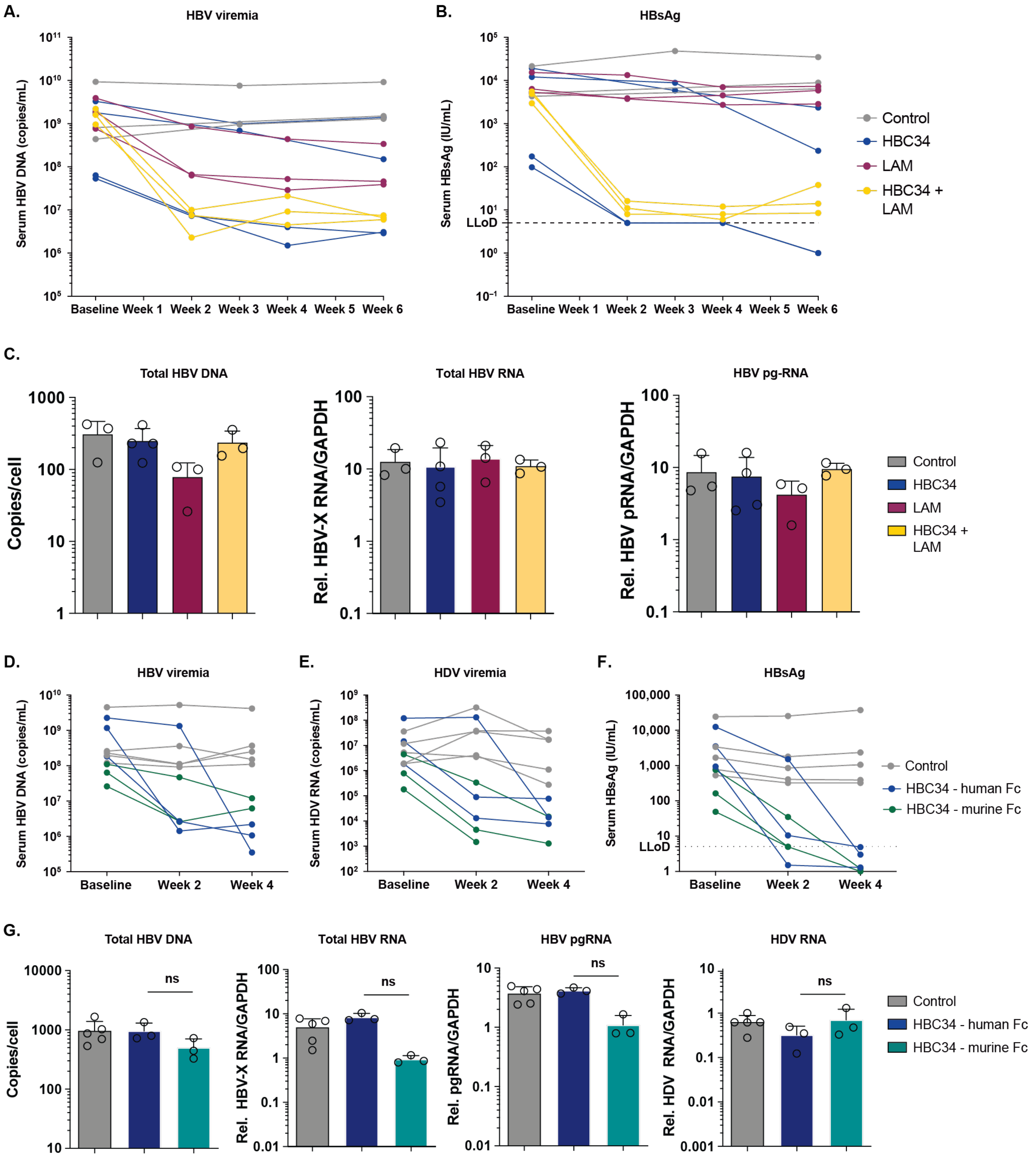

**SUPPLEMENTARY FIGURE S5. Antiviral treatment of chronically infected liver-chimeric mice.** Absolute levels of HBV viremia (A) and HBsAg (B) in HBV mono-infected liver-chimeric mice over the treatment period. (C) Mice were sacrificed at 6 weeks post treatment initiation and intrahepatic levels of HBV DNA, HBx RNA and pgRNA were quantified by (RT-)qPCR. Each circle represents one animal. Shown is the mean ± SD. Statistical differences relative to the control were analyzed by one-way ANOVA. p-value ns $p > 0.05$. (D-F) Absolute levels of HBV viremia (D), HDV viremia (E) and HBsAg (F) in HBV/HDV co-infected liver-chimeric mice over the treatment course. One animal in the murine Fc group was sacrificed at week 2. (G) Mice were sacrificed at 4 weeks post treatment initiation and intrahepatic levels of HBV DNA, HBx RNA, pgRNA and HDV RNA were quantified by (RT-)qPCR. Each circle represents one animal. Shown is the mean ± SD. Statistical differences relative to the control were analyzed by one-way ANOVA. p-value ** $p \leq 0.01$, ns $p > 0.05$.

**SUPPLEMENTARY TABLE S1. Full fit results for representative replicate of VIR-3434 binding to HBsAg by SPR**.

| HBsAg | VIR-3434 | ka1 (1/Ms) | kd1 (1/s) | KD1 (M) | ka2 (1/Ms) | kd2 (1/s) | KD2 (M) | Rmax1 (RU) | Rmax2 (RU) |
|---|---|---|---|---|---|---|---|---|---|
| Yeast | Fab | 2.64E+05 | 0.002062 | 7.80E-09 | 9.26E+05 | 1.14E-04 | 1.23E-10 | 13.39 | 12.3 |
| | IgG | 7.33E+05 | 8.83E-04 | 1.21E-09 | 4.56E+06 | 3.68E-08 | 8.06E-15 | 27.19 | 29.92 |
| PLC | Fab | 2.11E+05 | 1.39E-04 | 6.55E-10 | 9.77E+05 | 4.45E-04 | 4.55E-10 | 46.73 | 16.89 |
| | IgG | 2.52E+05 | 2.94E-04 | 1.17E-09 | 2.17E+06 | 3.94E-05 | 1.82E-11 | 82.9 | 85.63 |
